# The maternal-fetal interface of successful pregnancies and impact of fetal sex using single cell sequencing

**DOI:** 10.1101/641118

**Authors:** Tianyanxin Sun, Tania L. Gonzalez, Nan Deng, Rosemarie DiPentino, Ekaterina L. Clark, Bora Lee, Jie Tang, Yizhou Wang, Barry R. Stripp, Changfu Yao, Hsian-Rong Tseng, S. Ananth Karumanchi, Alexander F. Koeppel, Stephen D. Turner, Charles R. Farber, Stephen S. Rich, Erica T. Wang, John Williams, Margareta D. Pisarska

**Author notes:** Equally contributing authors (Tianyanxin Sun, Tania L. Gonzalez, Nan Deng).

## Abstract

The first trimester is a critical window of maternal-fetal communication for pregnancy. Therefore, we characterized crosstalk in ongoing human pregnancies at 11-13 weeks gestation. RNA-sequencing of matched maternal decidua and placenta identified 818 receptors and 3502 ligands, including 126 differentially expressed receptor-ligand pairs. Using single cell RNA-sequencing to further dissect placenta heterogeneity, we identified five major cell types (trophoblasts, stromal cells, hofbauer cells, antigen presenting cells and endothelial cells) with unique crosstalk at the maternal-fetal interface. We identified seven unique trophoblast subclusters, including new subtypes that transition into the terminal cell types, extra-villous trophoblasts and syncytiotrophoblasts. As fetal sex impacts pregnancy, we analyzed sex differences in each cell type and identified differences in immune cell function. TGFβ1, β-estradiol, and dihydrotestosterone emerge as upstream regulators of sexually dimorphic genes in a cell type specific manner. Thus, the fetal contribution at the maternal-fetal interface is cell and sex specific.

## Introduction

The human placenta is one of the most important organs in the body as it influences the health of mother and fetus during pregnancy. However, little is known about this organ in early pregnancy. This period, the first trimester is a critical window for placental development to ensure normal development of the fetus and maternal health throughout gestation. Aberrations in placentation can lead to adverse pregnancy outcomes throughout gestation such as miscarriages, small for gestational age, preeclampsia, placental abruption, placenta accreta and previa [1, 2]. Fetal sex can also impact these adverse outcomes, and we previously characterized the transcriptome profiles and sexual dimorphism of late first trimester placental tissue [3]. However, the placenta is a highly dynamic organ and consists of trophoblasts, mesenchymal and stromal fibroblasts, immune cells and other cell types that have unique functions for normal implantation and placentation. Therefore, it is critical to investigate single cell transcriptomic profiles in the first trimester placenta to better understand the dynamics of placental development. We set out to identify individual cell types, gene signatures as well as their enriched functions to characterize cell communication and the impact of fetal sex on placental development, with emphasis on trophoblast cells, since this population is most proximal to the maternal surface. A few recent studies systematically reported the single cell transcriptomes at the first trimester maternal-fetal interface from electively terminated pregnancies [4-6], but our study is the first to investigate single cell transcriptomes including cell type-specific sexual dimorphism in early placenta from ongoing healthy pregnancies that later resulted in live births. This will provide unique insights for future studies on cell type-specific and/or fetal sex-specific functions in early placental development that might impact maternal-fetal communication and contribute to overall outcomes.

## Results

### RNA-sequencing of matched decidua and placenta tissue from first trimester

To identify normal early gene expression at the maternal-fetal interface, total RNA-sequencing for matched decidua (maternal) and chorionic villi (fetal placenta) was performed. Principal component 1 shows that tissue type explains most of the difference between samples. Principal component 2 corresponds to fetal sex (**Figure S1**).

RNA-sequencing identified 16,035 genes reaching threshold FPKM>1 and Z-score>0 in decidua, placenta or both. Of these, 12,479 (77.8%) are protein-coding genes, including 3002 significantly differentially expressed genes (DEGs) with FDR<0.10 (**Figure S2A, Table S1**). Over a quarter of protein-coding DEGs are tissue-unique at the maternal-fetal interface, meaning they are expressed in either decidua or placenta, but not both. There are 2223 protein-coding DEGs upregulated in decidua compared to placenta (“decidua-upregulated”), including 827 only expressed in decidua (“decidua-unique”). There are 779 protein-coding DEGs upregulated in placenta compared to decidua (“placenta-upregulated”), including 60 only expressed in placenta tissue (“placenta-unique”).

### Receptors and ligands differentially expressed at the maternal-fetal interface

To investigate maternal-fetal communication, we focused on receptor and ligand DEGs. We identified 818 receptors expressed at the maternal-fetal interface (**TableS2**) [7-9] with 356 receptors significantly differentially expressed (FDR<0.10) between decidua and placenta, including 290 decidua-upregulated receptors and 66 placenta-upregulated receptors (**Figure S2B**). Half of decidua-upregulated receptors are also decidua-unique (146/290; 50.3%). The 5 decidua-unique receptors with highest fold-change over placenta are *PIGR* (FC = 215.3), *GABRP* (FC = 165.4), *CDHR1* (FC = 156.5), *TSPAN1* (FC = 149.1), and *SCARA5* (FC = 125.4) (**Table S2**). Of the placenta-upregulated receptors, 5 are placenta-unique: *NPFFR2* (FC over decidua = 3.18), *ADRA2B* (FC = 2.46), *OXER1* (FC = 2.20), *NRXN3* (FC = 1.99), and *ADGRL3* (FC = 1.99) (**Table S2**).

We identified 3502 candidate ligands, including extracellular or membrane-bound receptor-interacting protein-coding genes that do not fit a strict definition of “ligand” but will be referred to as ligands. There are 1043 ligand DEGs, including 773 decidua-upregulated and 270 placenta-upregulated ligands (**Table S3**). Similar to receptors, there are many decidua-unique ligands (266/773; 34.4%) and few placenta-unique ligands (17/270; 6.3%) (**Figure S2C**).

We identified 126 “DEG-DEG” receptor-ligand pairs with two genes significantly differentially expressed between decidua and placenta (**Figure 1A**). A subset of 27 receptor-ligands have potential maternal-fetal crosstalk, with one gene decidua-upregulated and the other placenta-upregulated (**Figure 1B**). Seven of these 27 include at least one gene with tissue-unique expression, including two placenta-unique genes (*CHAD* and *NRXN3*) and five decidua-unique genes (*CLCF1, CRLF1, POMC, SEMA3C*, and *TGFA*) (**Figure 1B**). The remaining 99 receptor-ligands are upregulated within the same tissue type, suggesting maternal or fetal self-signaling is strong in the first trimester (**Figure 1A, Table S4**). Most interactions (71/126; 56.3%) contained at least one gene that is tissue-unique (**Table S4**). Two hundred eleven “DEG-NS” receptor-ligands where one gene is significantly differentially expressed (FDR<0.10) and the other gene is not significant (NS) at the maternal-fetal interface were identified (**Figure S3**). Together, 337 receptor-ligands contain at least one gene that is significantly differentially expressed across the maternal-fetal interface (**Table S4**).

**Figure 1.**
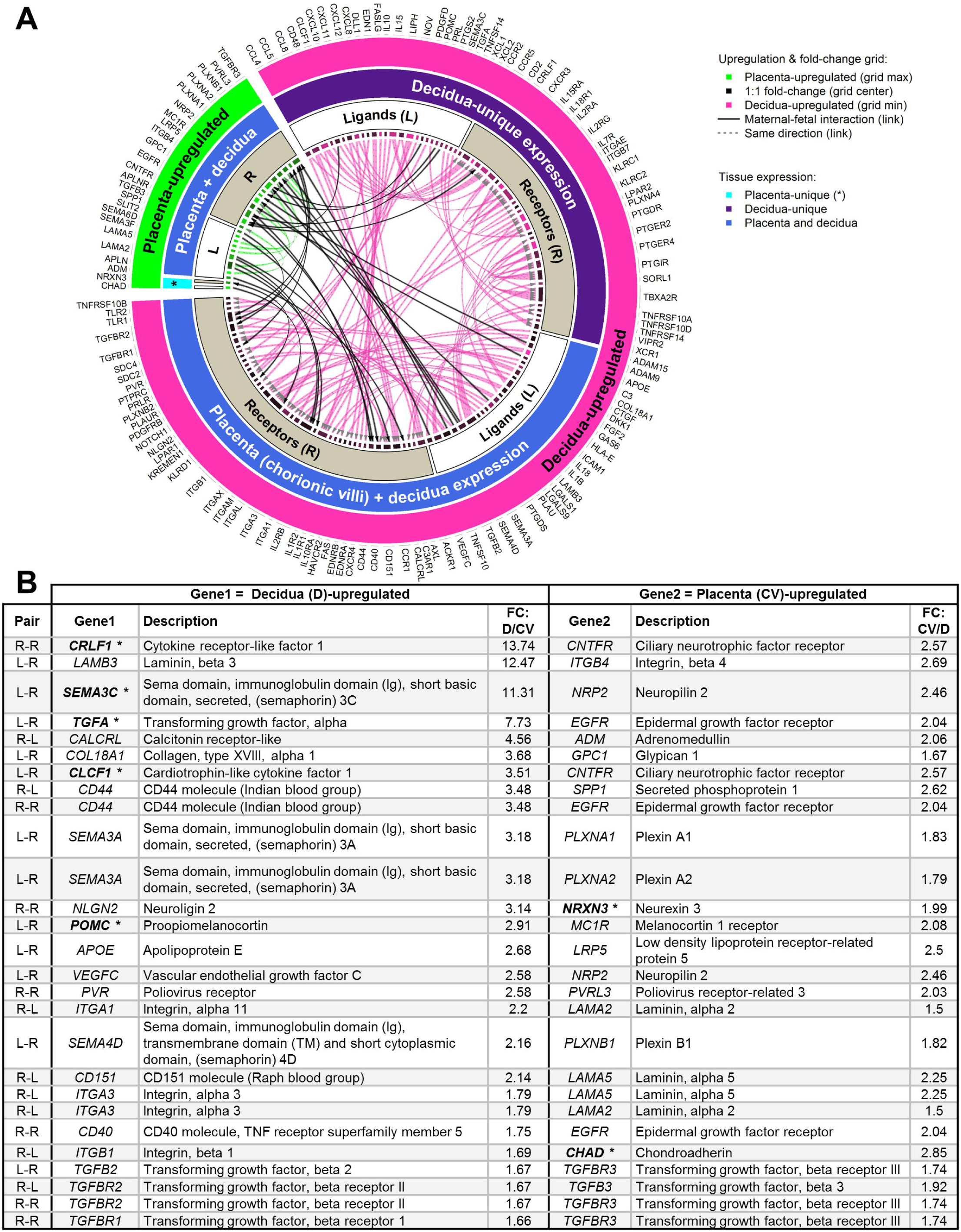
Receptor-binding interactions at the first trimester maternal-fetal interface. **(A)** Circos plot for 126 “DEG-DEG” gene pairs (both genes FDR<0.10). The innermost track is a fold-change grid showing magnitude and direction of upregulation. Arrowheads point to receptors. See **Table S4** for additional information such as FDR and FPKM values. **(B)** The 27 maternal-fetal interactions indicated by solid black links in the circos plot. Pair class: L=ligand, R=receptor. FC=fold-change. *Asterisk and bold: gene is also tissue-unique.

Ingenuity Pathway Analysis (IPA) identified decidua-upregulated receptors and ligands as highly enriched in immune and inflammatory functions, cell proliferation and migration, and vascular system functions (**Table 1**). Placenta-upregulated genes are associated with growth-promoting functions and pathways, including cell functions associated with other invasive tissue such as cancerous tumors, but predicted 2.867-fold decreased activation for “development of malignant tumors” demonstrating unique properties of placental invasion (**Table 1**).

**Table 1.** Enriched functions and disease associations of receptors and ligands differentially expressed at the maternal-fetal interface (top 30).

| Decidua-enriched functions and associations | Decidua P-value<sup>a</sup> | Placenta (CV)-enriched functions and associations | Placenta P-value<sup>a</sup> |
| --- | --- | --- | --- |
| Cell movement | 1.60E-115 | Adenocarcinoma | 1.92E-25 |
| Migration of cells | 4.20E-113 | Carcinoma | 3.77E-24 |
| Cell movement of blood cells | 3.00E-101 | Gastrointestinal adenocarcinoma | 4.83E-24 |
| Leukocyte migration | 7.20E-101 | Tumorigenesis of tissue | 9.21E-24 |
| Cell movement of leukocytes | 2.90E-99 | Intestinal adenocarcinoma | 3.12E-23 |
| Rheumatic Disease | 4.47E-92 | Large intestine adenocarcinoma | 8.85E-23 |
| Activation of cells | 1.02E-88 | Gastrointestinal carcinoma | 1.25E-22 |
| Inflammation of joint | 7.12E-88 | Non-melanoma solid tumor | 2.09E-22 |
| Systemic autoimmune syndrome | 5.16E-86 | Abdominal cancer | 2.74E-22 |
| Quantity of cells | 1.09E-85 | Gastrointestinal neoplasia | 8.16E-22 |
| Quantity of leukocytes | 2.39E-80 | Intestinal cancer | 8.84E-21 |
| Quantity of blood cells | 1.73E-78 | Cancer | 8.87E-21 |
| Cell movement of myeloid cells | 1.21E-75 | Gastrointestinal tract cancer | 1.04E-20 |
| Activation of blood cells | 4.27E-75 | Digestive system cancer | 1.21E-20 |
| Proliferation of blood cells | 5.53E-74 | Malignant neoplasm of large intestine | 2.19E-20 |
| Proliferation of lymphatic system cells | 9.14E-74 | Malignant solid tumor | 8.40E-20 |
| Cell movement of phagocytes | 1.34E-73 | Nonhematologic malignant neoplasm | 3.34E-19 |
| Cell death | 1.61E-73 | Abdominal neoplasm | 3.67E-19 |
| Activation of leukocytes | 2.32E-73 | Digestive organ tumor | 4.04E-19 |
| Development of vasculature | 2.67E-73 | Skin lesion | 4.04E-19 |
| Proliferation of immune cells | 3.38E-71 | Incidence of tumor | 1.60E-18 |
| Non-traumatic arthropathy | 1.20E-70 | Organismal death | 1.89E-18 |
| Inflammatory response | 6.18E-70 | Skin tumor | 3.50E-18 |
| Inflammation of organ | 1.52E-69 | Extracranial solid tumor | 6.65E-18 |
| Proliferation of lymphoid cells | 1.49E-68 | Cancer of secretory structure | 9.81E-18 |
| Angiogenesis | 2.55E-68 | Skin cancer | 1.00E-17 |
| Proliferation of lymphocytes | 6.93E-68 | Cell movement | 1.57E-17 |
| Interaction of blood cells | 7.19E-68 | Frequency of tumor | 1.67E-17 |
| Binding of blood cells | 1.22E-67 | Development of malignant tumor | 1.89E-17 |
| Proliferation of mononuclear leukocytes | 1.59E-67 | Genitourinary tumor | 2.15E-17 |

<sup>a</sup> P-value = Fisher's Exact Test.

### Single-cell transcriptomic profiling

Since the placenta consists of different cell types, we dissected the heterogeneity of first trimester placental cells using single cell RNA-sequencing. To maximize the cell populations captured, cells were collected without marker selection. Five clusters (P1-P5) were separated from unsupervised clustering **(Figure 2A)**. We identified clusters by expression of established markers: trophoblast cells (*KRT7, KRT8, CGA, EGFR*), stromal fibroblast cells (*COL3A1, PDGFRA, THY1*), hofbauer cells (*CD14, CD163, CSF1R*), antigen presenting cells (APCs) (*HLA-DRA, HLA-DPB1, CD52*) and endothelial cells (*PECAM1, CD93*) **(Figure 2B)**. Stromal fibroblast cells (P2) are the most abundant cell type (38.8%), followed by hofbauer cells (P3, 29.4%), trophoblast cells (P1, 22.1%), APCs (P4, 8.5%), and endothelial cells (P5, 1.2%). DEG analysis and pairwise comparison yielded genes that are specific to each cluster corroborating the identification of each cluster **(Figure 2C, Table S5)**.

**Figure 2.**
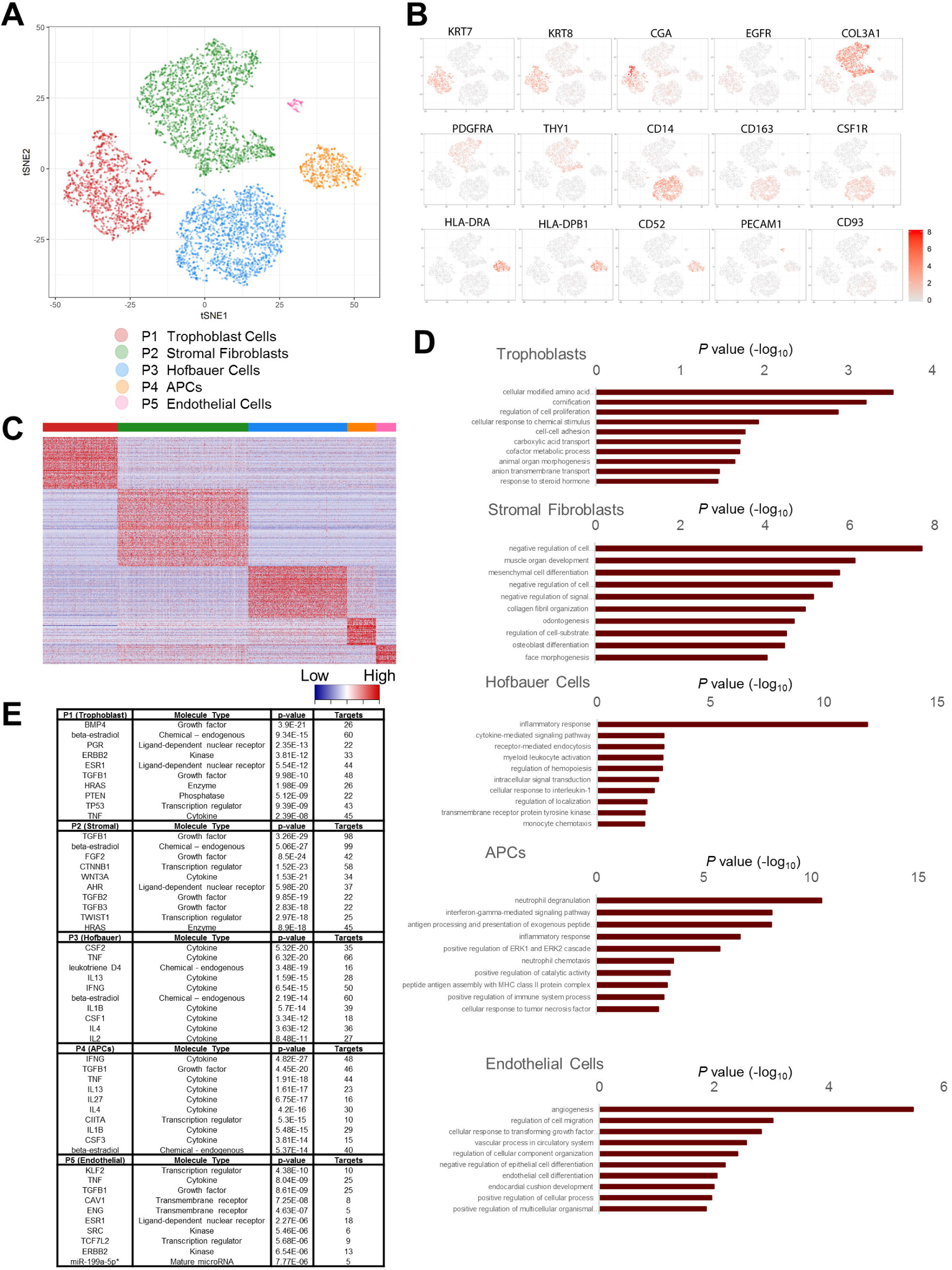
Single cell transcriptomic profiles of the first trimester placenta. **(A)** Placental cell clusters visualized by t-Distributed Stochastic Neighbor Embedding (t-SNE) using Scan and Scater packages. Colors indicate cell types. **(B)** Gene expression of selected markers used to distinguish the five major clusters in tSNE plot. Log-transformed, normalized expression levels are shown. **(C)** Heatmap of differentially expressed genes from pairwise comparison, for the five major clusters shown by cells (columns) and genes (rows) with the expression level noted. Color bar indicates the cell types. **(D)** The top 10 enriched GO terms using cell type specific markers are presented as −log_10_(P value) for each cell type, Bonferroni-corrected for P < 0.05. **(E)** The 10 most significant upstream regulators predicted by IPA are shown for each cell cluster. Input is the significantly and specifically expressed genes in each cell type using DEG analysis and pairwise comparison. APCs: antigen presenting cells. * includes other miRNAs w/seed CCAGUGU.

Trophoblasts (P1) specifically express the epithelial cell keratin genes (*KRT7, 8, 18, 19, 23*) and cancer testis antigen genes (*PAGE4, XAGE2/3, MAGEA4*) [10]. The stromal fibroblast population contains cells at different stages of differentiation, including undifferentiated mesenchymal stem cells and differentiated fibroblasts as well as myofibroblasts. Mesenchymal stem cell marker *NT5E* and stem cell markers *NANOG, OCT4* [11, 12] are not captured in our cell population but the cytoskeletal filament genes *DES* and *ACTA2*, markers for fibroblasts and myofibroblasts, are both highly and specifically expressed in P2, suggesting that the stromal fibroblast population from dissociated placental samples are primarily in their differentiated state. Extracellular matrix regulatory genes are also specifically expressed in the stromal fibroblast population, including 17 collagen encoding genes such as *COL1A1*, non-collagenous glycoproteins such as *LAMA2, LAMC3*, proteoglycans such as *DCN* and matrix metalloproteinases such as *TIMP1* and *TIMP3*. P3 and P4 are characterized as 2 immune populations. They abundantly express myeloid lineage markers *AIF1* and *CD53*, lack *CD56* (marker of natural killer cells) and express chemokine receptors (*CXCR4, CCR1, CCR7*) that natural killer cells normally do not express [13], and therefore are likely macrophages or dendritic cells. Unique to the P3 population is expression of *CD14* and *CSF1R* suggesting that P3 consists of a macrophage population [14]. Although both maternal and fetal macrophages reside within the first trimester placentas [4] and are important in phagocytosis for innate immunity [15] and tissue remodeling [16], the P3 population expresses *CD86, LYVE1* and lacks expression of MHC type II antigens, *HLA-DR*, -*DP*, and -*DQ*, consistent with first trimester fetal hofbauer cells [17] and devoid of maternal macrophages that are known to express *HLA-DR* with low *CD86* expression [18]. Hofbauer cells arise from mesenchymal stem cells in early placenta and are supplemented from fetal bone marrow-derived macrophages once fetal circulation is established [19]. Hofbauer cells are heterogenous but characterized by high expression of *CD163* (hemoglobin scavenger receptor, specifically expressed by P3) [20] and represent an active line of fetal defense against vertical viral transmission and maternal IgG antibodies, by presenting Fc receptors (*FCGRT/1A/1B/2A/P*, specifically expressed by P3) [21] as well as phagocytotic properties. Hofbauer cells also play a role in villous branching morphogenesis (*SPRY2*, specifically expressed by P3), placental vasculogenesis (*LYVE1*, specifically expressed by P3, and *VEGFB*, although not specifically expressed by P3, is the highest in P3) as well as synthesizing cytokines (*CCL2, CCL3, CCL4, CCL13, CCL4L2, CCL3L3*, specifically expressed by P3). The P3 population also specifically expresses hemoglobin genes such as *HBA1, HBA2, HBG2*, supporting the hematopoietic origin of hofbauer cells and their role in ingestion of nuclei of primitive red blood cells in the first trimester placental villi [22]. Alternatively, P4 specifically expresses a group of MHC class II antigens (or regulatory) molecules *HLA-DR, HLA-DP, HLA-DQ, HLA-DM* as well as *CD52, CXCR4, SERPINA1, MMP7, MMP9* suggestive of professional APCs such as dendritic cells and/or macrophages. P5 is the least abundant cell population and specifically expresses *PECAM1* and *CD93*, markers of endothelial cells.

### Enrichment analyses in first trimester placenta cells

Gene ontology enrichment analysis with cell type-specific genes identified enriched biological processes **(Figure 2D)**. The top enriched processes in trophoblasts include metabolic processes of amino acids and cofactors, regulation of cell proliferation, cell adhesion, cargo transport and response to steroid hormones, indicating highly active metabolism and morphological dynamics [23], and unique functions in mediating hormonal signals and exchange of metabolic products between maternal and fetal compartments. Stromal fibroblasts are enriched in cell-substrate adhesion and negative regulation of cell proliferation, differentiation and signal transduction. In both hofbauer cells and APCs, inflammatory responses and signaling pathways mediated through cytokines are among the top enriched terms, highlighting their essential roles in conducting immune functions. Endothelial cells are enriched in cell differentiation, migration and angiogenesis, consistent with development of the vasculature system in a healthy first trimester placenta.

### Cell type-specific upstream regulators

*TGFB1* and β-estradiol were identified as significant upstream regulators of all cell types (*TGFB1* for hofbauer cells and β-estradiol for endothelial cells are significant, though not top 10) **(Figure 2E)**. The most important upstream regulators for trophoblast cells are growth factor *BMP4*, hormone nuclear receptors *PGR* and *ESR1*, kinase *ERBB2*, enzyme *HRAS*, phosphatase *PTEN*, transcription regulator *TP53* and cytokine *TNF*. In addition to *TGFB1* and β-estradiol, the top regulators for stromal fibroblasts are growth factors *FGF2, TGFB2, TGFB3*, transcription regulators *CTNNB1* and *TWIST1*, nuclear receptor *AHR*, cytokine *WNT3A* and enzyme *HRAS*. In contrast, immune cell populations are regulated mainly by cytokine members, such as *CSF1/2, TNF, IL13, IFNG, IL1B, IL4 and IL2* for hofbauer cells, and *IFNG, TNF, IL13, IL27, IL4, IL1B* and *CSF3* for APCs. The endogenous chemical leukotriene D4 and transcription regulator *CIITA* are also upstream regulators for hofbauer cells and APCs, respectively.

### Upstream regulators, crosstalk between maternal decidua and fetal placental cells

Of the upstream regulators, the majority are not decidua-or placenta-unique, suggesting signaling to the various cell types may be maternal-fetal, fetal-fetal or both. Hormones, β-estradiol and progesterone are produced by STBs (differentiated trophoblast cells), and their respective receptors, *ESR1* and *PGR*, are decidua-unique, demonstrating maternal-fetal crosstalk at the interface which regulates trophoblast cell function. *TGFB1* is also an upstream regulator expressed in both decidua and placenta, as are its receptors, *TGFBR1* and *TGFBR2*. Stromal fibroblast regulation is impacted by decidua-upregulated fibroblast growth factor 2 (*FGF2*) through its receptor *SDC2* (syndecan 2, decidua-upregulated) and receptor *SDC4* (syndecan 4, decidua-upregulated). Stromal fibroblasts are also regulated by another *TGFB* family member, placenta-upregulated *TGFB3*. Among the cytokines regulating the immune populations, decidua-upregulated interleukin 1β (*IL1B*) is an upstream regulator of both hofbauer cells and APCs.

### Cell unique crosstalk

We investigated how different placental cell types coordinate communication with the maternal decidua and amongst themselves through receptor-ligand interactions. Genes specifically expressed in each of the five identified cell clusters were matched with interacting gene partners from **Table S4** to identify crosstalk. We identified 58 receptor-ligand pairs between the placental cell markers and decidua-upregulated genes, including 27 interactions with decidua-unique genes (**Figure 3A, 3B**). Among these interactions, a few maternal-fetal receptor-ligand pairs from tissue RNA-seq had cell specificity. This includes the decidua-unique genes cytokine receptor-like factor 1 (*CRLF1*) and cardiotrophin-like cytokine factor 1 (*CLCF1*), which interact with the stromal fibroblast-specific marker ciliary neurotrophic factor receptor (*CNTFR*) (**Figure 1B, 3A, 3B**). The differentially expressed decidua-placenta pair *CD44-SPP1* is an APC-hofbauer interaction in single cell analysis.

**Figure 3.**
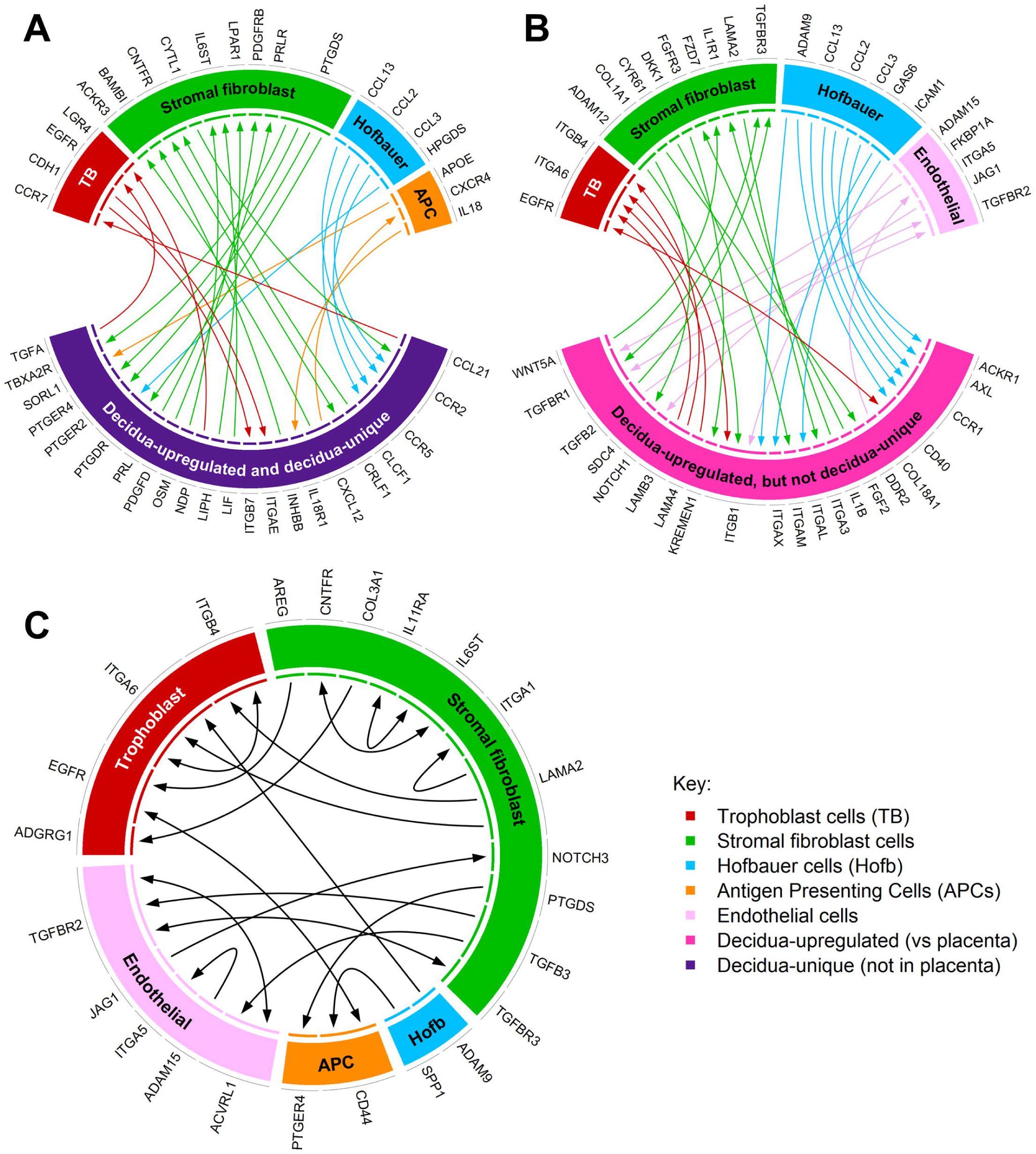
Circos plots with placental cells identified by single cell RNA-sequencing. **(A)** Interactions between placental cell markers (single cell RNA-seq) and decidua-unique genes (total RNA-seq). **(B)** Interactions between placental cell markers (single cell RNA-seq) and decidua-upregulated (not unique) genes (total RNA-seq). **(C)** Interactions among cell markers for trophoblast, stromal fibroblast, hofbauer, APCs (antigen presenting cells), and endothelial cells.

Most placental cell clusters expressed cell-surface marker genes with binding partners encoded by decidua-unique genes, except the smallest cluster representing endothelial cells. We also highlighted 18 gene pairs (comprised of 24 cell markers) involved in crosstalk between placental cell clusters (**Figure 3C**), indicating placental cell-cell communication.

### Heterogeneity and developmental reconstruction of trophoblasts

A total of 1465 trophoblast cells were further dissected into sub-clusters T1∼T7 (constituting 41.6%, 19.1%, 9.5%, 9.3%, 10%, 6.5% and 4% of the trophoblast population, **Figure 4A**). Cytotrophoblast markers are expressed throughout different subclusters. *PARP1* is abundantly expressed in T1, T2, T3, and T4. *ITGA6* is abundantly expressed in T4 and T6. The expression of *TP63* is greatest in T4. The expression of a recently reported cytotrophoblast marker *NRP2* [4] is highest in T3 and T4. Conventional markers and differential gene expression analysis characterized T5 and T6 to be STBs and EVTs, respectively (**Figure 4B, Table S6**). T5 specifically expresses retroviral envelope genes *ERVFRD-1* and *ERVV-1* that are critical for cell fusion, hormone synthesis genes *CYP19A1* (converts androgens to estrogens) and *HSD11B2* (converts cortisol to cortisone), as well as solute carrier SLC family members such as *SLC40A1* and *SLC26A2*. T6 specifically expressed genes important for cell adhesion, cell invasion, extracellular remodeling (*TIMP3, SPON2, MFAP5*) and immunomodulation (*TGFB1, HLA-G*). Consistent with the highly proliferative activity of the cytotrophoblasts in the first trimester placenta, a group of cell cycle regulators were specifically expressed in T2 (*MKI67, CCNA2, CCNB1/2, CDK1*), T4 (*CDK6*), and T6 (*CDK7* and *CCNE1*). Cytotrophoblasts consist of multiple sub-clusters and we reconstructed the developmental trajectory **(Figure 4C)** that demonstrated a bifurcating pattern of trophoblast cell development. One branch ends with EVTs (T6) and the other ends with STBs (T5). T2 and T4 are the initial clusters and likely the stem cytotrophoblasts. T1, T3 and T7 occupy the intermediate stages, suggesting they are the transitional stages leading to the fully differentiated EVTs or STBs.

**Figure 4.**
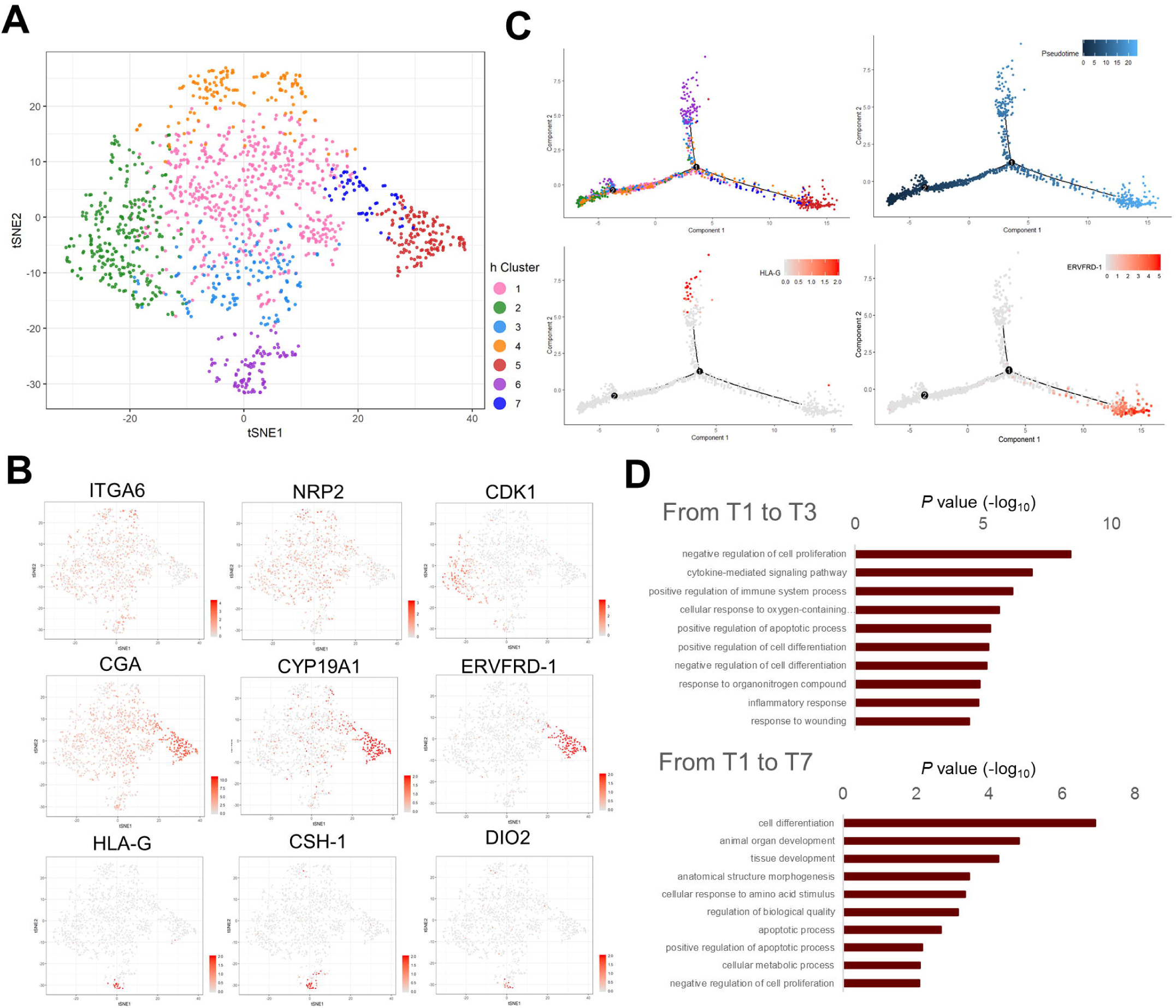
Single cell transcriptomic profiles of trophoblast cells within the first trimester placenta. **(A)** tSNE of trophoblast sub-clusters using Scan and Scater packages. Colors indicate sub-clusters within the trophoblast population. **(B)** Gene expression of selected markers used to distinguish selected sub-clusters Log-transformed, normalized expression levels are shown. **(C)** Pseudotime reconstruction of the developmental trajectory in the trophoblast population shows trophoblast subclusters (T1∼T7), indicated by colors. The timepoints 1 and 2 represent hierarchical branching events. Pseudotemporal trajectory indicates developmental timeline. The distribution patterns of cells expressing EVT marker gene *HLA-G* and STB marker gene *ERVFRD-1* in the pseudotemporal trajectory. **(D)** The top 10 enriched GO terms are presented as −log_10_(P value) for each cell developmental progression. (Bonferroni-corrected for P < 0.05).

GO term enrichment analyses to understand the transitional stages and T1 to T3 (EVT path) revealed immune related functions such as cytokine-mediated signaling pathways, positive regulation of immune system processes and inflammatory responses as well as regulation of cell proliferation, differentiation and apoptosis **(Figure 4D).** Enriched GO terms from T1 to T7 (STB path) revealed increased metabolic processes and cellular response such as that to amino acid stimulus. The regulation of cell proliferation, differentiation and apoptosis was also emphasized in this path, especially positive regulation of apoptotic process and negative regulation of cell proliferation, consistent with the known properties of STB, which undergo constant turnover [24] but are still metabolically active [25].

### Sexual dimorphism in the placenta

We previously characterized the first trimester placenta transcriptome at the tissue level and identified sexually dimorphic genes that may contribute to different pregnancy outcomes among sexes [3]. Sex differences are also present at the global gene expression level in decidua and placenta **(Figure S1)**. Therefore, we investigated the sexual dimorphism at the single cell level in the first trimester placenta. Of the 7,245 cells that are selected for analysis, 4,369 cells are from the female placentas and 2,876 cells are from the male placentas **(Figure 5A)**. The top genes (Ilog_2_FCI>1, FDR<0.01) significantly upregulated in cells from either sex are shown in **Figure 5B**, among which genes that are also specific markers to corresponding cell types are of particular interest, including *MUC15, NOTUM* and *MAGEA4* in trophoblasts, *STC1* in stromal fibroblasts, *FCGBP, CCL13* and *RETN* in hofbauer cells and *IL1RN, MMP9 and GPR183* in APCs. The fold changes of top DEGs among sexes are shown in **Table S7**.

**Figure 5.**
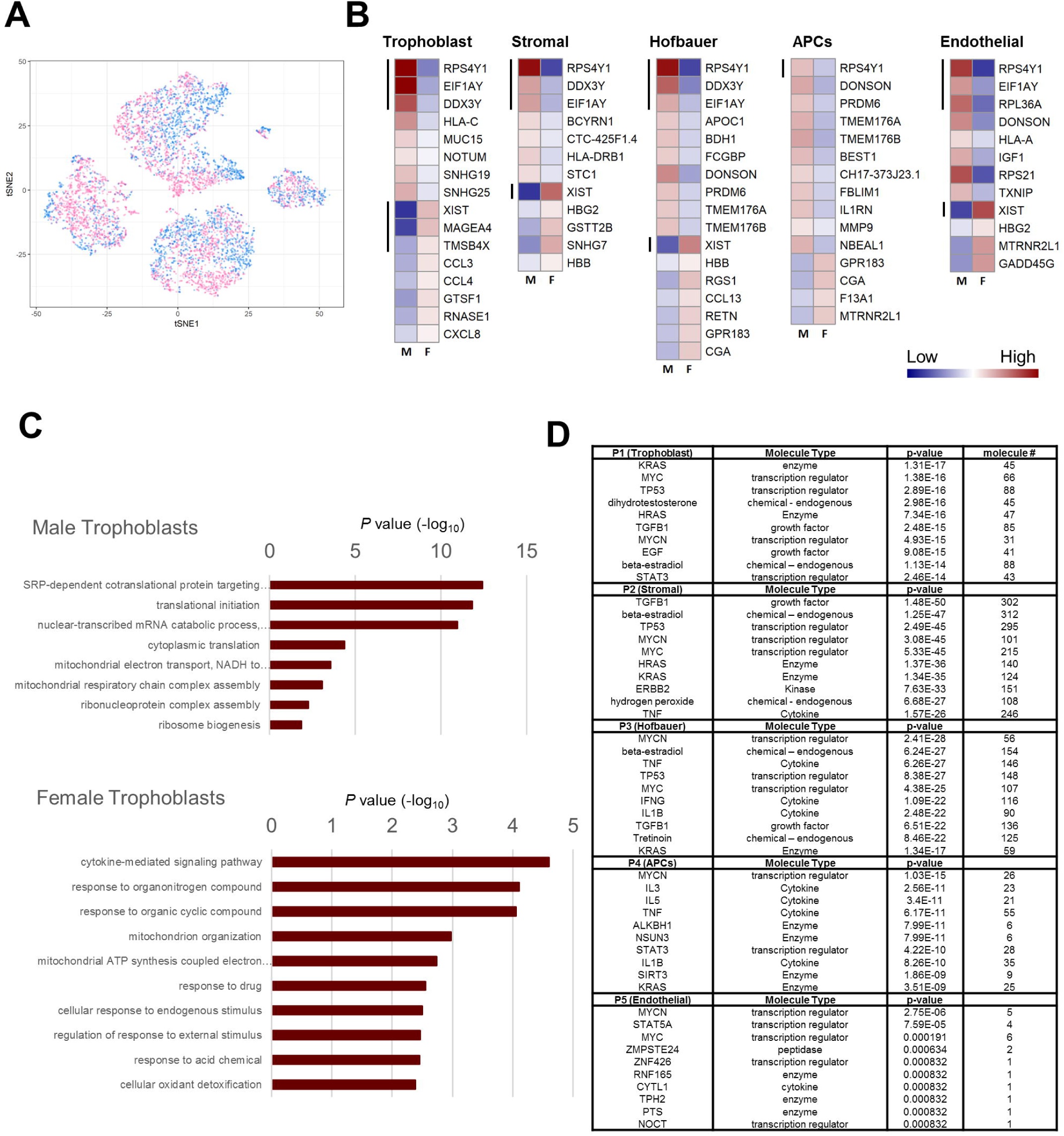
Sexual dimorphism of single cell transcriptomic profiles in the first trimester placenta. **(A)** tSNE of male and female placenta cells with males colored in blue and females in pink. **(B)** The top sexually dimorphic genes (FDR<0.01, ILog_2_FCI>1) in each cluster are presented as a heatmap. Black bars on the left of the heatmap indicate genes on sex chromosomes. **(C)** The top 10 enriched GO terms are presented as −log_10_(P value) for either sex in every cell type. (Bonferroni-corrected for P < 0.05). **(D)** The 10 most significant upstream regulators of sexually dimorphic genes predicted by IPA are shown using the differentially expressed genes between sexes for each cell cluster. APCs: antigen presenting cells.

Essential for translational initiation and RNA modification, three genes on the Y chromosome (*DDX3Y, EIF1AY, RPS4Y1*) are upregulated in almost all cell types of male placentas compared to female placentas, except for *DDX3Y* and *EIF1AY* in APCs, and *DDX3Y* in endothelial cells. *RPL36A* (a ribosomal gene on X chromosome) is only upregulated in endothelial cells of male placentas. Only trophoblasts cells have unique X chromosome genes upregulated in females compared to males including *MAGEA4* (melanoma associated antigen 4) and *TMSB4X* (thymosin beta 4) whereas *XIST* (X inactive specific transcript) is upregulated in all cell types except APCs, which are likely maternally derived.

The majority of the DEGs among sexes are autosomal genes, with 27 significantly upregulated in males and 17 significantly upregulated in females (Ilog_2_FCI>1, FDR<0.01). These genes are involved in cell-cell interaction, antigen presenting, small regulatory RNAs and metabolic functions. In trophoblasts, *HLA-C, MUC15, NOTUM, SNHG19* and *SNHG25* are upregulated in males. *BCYRN1, CTC-425F1.4, HLA-DRB1* and *STC1* are upregulated in male stromal fibroblasts. *APOC1, BDH1, FCGBP* are upregulated in male hofbauer cells and *BEST1, CH17-373J23.1, FBLIM1, IL1RN, MMP9* and *NBEAL1* are upregulated in male APCs. Male endothelial cells have upregulation of *HLA-A* and *IGF1* compared to females. Several genes are upregulated in males in more than one cell type which includes *DONSON*, expressed in hofbauer cells, APCs, and endothelial cells, as well as *MT1G, PRDM6, TMEM176A* and *TMEM176B* upregulated in male hofbauer cells and APCs. Similarly, of the 17 genes upregulated in female placentas, some are uniquely upregulated in individual cell types of female placentas while others are shared between cell types. *CCL3, CCL4, GTSF1, RNASE1* and *CXCL8* are upregulated in female trophoblasts. *GSTT2B* and *SNHG7* are upregulated in female stromal cells. In hofbauer cells, *RGS1, CCL13* and *RETN* are upregulated in females and *F13A1* is upregulated in APCs. *GADD45G* is specifically upregulated in female endothelial cells. Genes that are upregulated in females in multiple cell types include *HBG2* in stromal fibroblast cells and endothelial cells, *HBB* in stromal fibroblast cells and hofbauer cells, *GPR183* and *CGA* in hofbauer cells and APCs as well as *MTRNR2L1* in APCs and endothelial cells.

Enrichment analyses revealed enriched GO terms in male and female placentas **(Figure 5C, Figure S4)**. Male trophoblast cells are enriched in protein translation, certain mitochondrial and ribosomal functions whereas female trophoblast cells are mostly enriched in responses to various compounds and stimuli and cytokine-mediated signaling pathway. Stromal fibroblast cells from both fetal sexes are enriched with signal recognition particle-dependent co-translational protein targeting to membrane, translational initiation and nuclear-transcribed mRNA catabolic process, indicating the importance of these conserved functions between sexes as well as the diversity of genes from either sex that contribute to these fundamental processes. Furthermore, regulation of cell migration, angiogenesis and cellular component organization are enriched in male stromal fibroblast cells whereas in female stromal fibroblast cells, regulation of apoptotic processes, viral processes and neutrophil degranulation are enriched. Male and female hofbauer cells also shared several enriched terms. Specifically, male hofbauer cells are enriched in cell surface receptor signaling, regulation of cell differentiation and cellular responses to stress and cytokine stimuli, while female hofbauer cells are enriched in cellular responses to chemical and organic stimuli and negative regulation of apoptotic process. Within the APC population, sex differences include enriched response to oxidative stress, cytokine-mediated signaling pathway, positive regulation of T cell activation and negative regulation of apoptotic process in female cells. In the endothelial population, male cells are enriched in translational regulation and ribosomal biogenesis.

To understand what regulates the DEGs between fetal sexes, we performed upstream regulator analyses using IPA (top 10 regulators are shown in **Figure 5D**). Transcriptional regulators *MYCN* and/or *MYC* are upstream regulators for sexually dimorphic genes in all five cell types. TGFβ1 and β-estradiol, the top significant upstream regulators found in most clusters (**Figure 2E**), also regulate sexually dimorphic genes in trophoblasts, stromal fibroblast cells and hofbauer cells (**Figure 5D)**. In addition, the hormone, dihydrotestosterone impacts sexually dimorphic genes in the trophoblast population. Sex differences in hofbauer cells and APCs, the two immune populations, are also regulated by cytokines such as *TNF* and the interleukin family.

## Discussion

This is the first transcriptomic study of the human first trimester maternal-fetal interface in healthy ongoing pregnancies, ensuring that the transcriptomes are indicative of normal placentation and uncomplicated human pregnancies. Recently, human placental transcriptomes and interactions at the maternal-fetal interface were described at the single cell level. Initial studies were performed using human term placenta [26, 27], reflective of terminal placental functions which differ from early placentation [6]. Early in gestation, identification of placental cell types and maternal-fetal communication have only been performed following terminations, which may not reflect healthy ongoing gestations as pathology may be present leading to the loss of the pregnancy [4, 5]. Single cell studies of placental tissue have also been performed using murine tissue [28], however lack of decidual tissue in this data set makes it less useful to study the maternal-fetal interface.

In addition to the identification of the normal transcriptome at the maternal-fetal interface, this study is unique in the identification of the first trimester single cell transcriptome of ongoing pregnancies without marker selection, thus providing a more unbiased identification of signature transcripts of previously described cell types [4, 6]. We identified APCs within the placenta which exhibit dendritic-like properties, which may be a distinct feature of ongoing healthy pregnancies, demonstrating the significance of the interwoven immune regulation between mother and fetus at the interface.

Maternal decidua signaling is highly enriched in immune and inflammatory functions, cell proliferation and migration, and vascular system functions whereas placentas are strongly associated with growth-promoting functions and pathways. The 3-way cytokine-signaling interactions between *CNTFR* (placenta-upregulated), C*RLF1* and *CLCF1* (both decidua-unique) [29], suggests cytokine mediated signaling that is both fetal-maternal and maternal-maternal. *CNTFR* upregulation promotes cell proliferation and potential cell invasion in human gliomas [30], thus these maternal-fetal interactions may indicate immunological signaling impacting placental cell proliferation and invasion for a successful pregnancy. Maternal-fetal interactions also include the extracellular matrix, such as decidua-upregulated *LAMB3* which binds placenta-upregulated *ITGB4*, both genes that promote cell proliferation, migration, and invasion [31, 32], all necessary processes for trophoblast cells to establish a normal placenta. We identified *ITGB4* as a unique marker of trophoblast cells. *LAMB3*, which plays an important role in the endometrial extracellular matrix during early pregnancy [33], may be signaling trophoblasts through *ITGB4* and mediating invasion into maternal tissue during placentation, and needs to be studied further.

*TGFB1*, expressed in decidua and placenta in total RNA-seq, is a key upstream regulator in all cell types, confirming TGFβ1 is a significant coordinator of placental development at the maternal-fetal interface [34]. *TGFB1* is specifically expressed in the EVT subcluster in trophoblasts which may coordinate invasion of EVTs with differentiation of STBs through its receptor *TGFBR1*, which is expressed in both EVT and STB subclusters in our study (data not shown) and by others [27]. Another TGFβ superfamily member, BMP4, is a key upstream regulator in the trophoblast population. BMP4 is critical for initiating differentiation of human embryonic stem cells to trophoblasts [35] and for studying human trophoblast lineages derived from human pluripotent stem cells [36].

Precise hormonal regulation is necessary for a healthy gestation, regulated through maternal-fetal crosstalk initiated by the mother and later by the fetus through the placenta. Estrogen and progesterone production switches from the luteal cells in the maternal ovary to the STBs in the placenta at 7∼9 weeks gestation, which is fully established by 10 weeks, making the placenta the source of hormone production essential for pregnancy maintenance [37]. β-estradiol is a top key upstream regulator in all placental cell types identified from placentas at 11-13 weeks gestation in our studies. Receptors for β-estradiol (*ESR1*) and progesterone (*PGR*), both decidua-unique, are among the top upstream regulators in trophoblast cells, suggesting at this gestational age, signaling to the maternal decidua is from hormones produced by the STBs, which through crosstalk regulate the placenta and trophoblasts. The critical role of precise hormonal regulation in placentation has been identified during aberrant hormonal states, including supraphysiologic hormone states which lead to placental reprogramming resulting in elevated estradiol and progesterone production. Elevated hormone production affects transcriptional regulators of trophoblasts, such as *GATA3*, altering trophoblast migration and invasion, which can lead to placental dysfunction and adverse outcomes including low birth weight and small for gestational age (SGA) infants [2, 38-41].

The majority of cell cluster upstream regulators also regulated sexually dimorphic genes within each cluster, with 54.7% (276/505) of regulators for trophoblasts also being regulators of sexually dimorphic genes in trophoblasts, similarly for stromal cells (66.5%; 647/973) and hofbauer cells (61.7%; 539/874). However, only 33.6% of regulators in APCs are also regulators of sexually dimorphic genes, which may be maternally derived and less directly influenced by fetal sex. The steroid hormone DHT (5α-dihydrotestosterone) is a unique upstream regulator of male trophoblasts. Testosterone is produced in the fetal testis at 8 weeks gestation [42], suggesting there is a significant contribution of the fetus in placental regulation. Testosterone in humans may impact placental differentiation similar to other animal models [43, 44] which produce sexually dimorphic outcomes attributed to placental function, such as fetal growth [45].

Marker genes were identified to be sexually dimorphic. Among the trophoblast population, X-linked *MAGEA4*, is upregulated in female trophoblasts, and autosomal *MUC15* and *NOTUM*, are upregulated in male trophoblasts. *MAGEA4* is also the most significantly placenta-upregulated DEG compared to decidua in our total RNA-seq (FDR=1.63 × 10^-15^). *MAGEA4* is an oncogene associated with tumor invasiveness and aggressiveness [46]. *NOTUM* is a polarity determinant that affects stem cell migration [47] and *MUC15* is important for cell adhesion to the extracellular matrix [48]. Collectively, these genes may play significant roles in sexually dimorphic trophoblast migration and invasion providing insight into sexually dimorphic placental dysfunction and disease, such as preeclampsia [48].

We identified 7 unique subclusters of trophoblast cells with new subtypes that transition into the terminal cell types, EVTs and STBs. The GO terms of the transitional processes identified important functional dimorphism between the EVT path and STB path. Immune related functions and increased metabolic processes are highlighted for the EVT path and STB path, respectively. *HLA-G*, the classical cell type-specific marker for EVT cells [49], is only detected in a portion of the EVT population, whereas the rest of EVTs identified by hierarchical clustering and semi-supervised developmental reconstruction lack *HLA-G* expression. *HLA-G+* cells are located at the endpoint of the pseudotime axis and therefore are truly terminally differentiated EVTs while these other EVTs may be transitional states.

There are 2 immune populations, hofbauer cells and APCs identified in the first trimester placenta. Hofbauer cells are villous macrophages of fetal origin that functionally resemble M2 macrophages which play important roles in placental vasculogenesis and angiogenesis. The expression of *ITGAX* (*CD11c*) and mature dendritic cell marker *CD83*, together with a lack of *DC-SIGN* and moderately expressed *CD14* (both are immature dendritic cell markers) suggests that the APC cluster contains a dendritic population of varying states of maturation [14, 50]. Further studies are necessary to characterize the heterogeneity and subtypes within the APC population from first trimester ongoing pregnancies. The decidua-placenta interacting pair, *CD44-SPP1* is also an APC-hofbauer interaction, which independently has been identified to play roles in migration and invasion in cancer [51, 52] as well as at the maternal-fetal interface [53]. Based on our findings, controlled migration and invasion may be due to immunologic regulation between the maternal-derived APCs and the fetal-derived hofbauer cells.

Immunologic regulation appears to also be sexually dimorphic, as a number of genes upregulated in female cells are chemokines such as CCL3 in trophoblasts and CCL13 in hofbauer cells. This strongly suggests that certain immune functions are enhanced or altered in female placentas compared to male placentas. CCL3, through decidua-upregulated CCR1, is involved in recruitment of natural killer cells and monocyte migration from the maternal interface [54, 55]. *CCL13*, with higher expression in female hofbauer cells, binds to decidua-unique *CCR2*, and functions as a chemoattractant for various immune cells, suggesting that hofbauer signaling for immune cell migration is sexually dimorphic, impacting the maternal immune response differently. In addition, *CCL13* is involved in the M2 phenotype of macrophages [56]. Increased expression of *CCL13* in females may increase the M2 phenotype, which mainly induces the Th2 response and has greater involvement in tissue repair than mounting a maternal response, consistent with the premise that male conceptions promote increased maternal innate and adaptive immunity compared to female conceptions [57]. This is also highlighted in the enriched responses to oxidative stress, cytokine-mediated signaling pathway, positive regulation of T cell activation and negative regulation of apoptotic process in maternal derived APCs from female gestations.

In summary, the maternal-fetal interface in ongoing human pregnancies is a hormonally and immunologically controlled environment for the establishment and development of the placenta. Hormonal regulation, initially from the maternal ovarian luteal cells followed by subsequent regulation from the fetal STBs, maintains the maternal surface to immunologically balance the invasion of the fetal trophoblasts that are critical for the establishment of the placenta, through maternal-fetal crosstalk. In addition, the TGFβ family, specifically TGFβ1, is critical for maternal-fetal communication and coordination of trophoblast differentiation to ensure appropriate timing of EVT invasion with STB development, important for nutrient and gas exchange and hormone production. Through single cell sequencing we identified genes that are sexually dimorphic, from both the sex chromosomes and autosomes that may undergo different functions in a cell type-specific manner, with female trophoblasts more likely to be enriched in the cytokine-mediated signaling pathway. In addition to overall regulation, TGFβ1 and β-estradiol are also identified as significant upstream regulators of sexually dimorphic genes including hofbauer cells, suggesting immune mediated pathways are important in female gestations. As maternal decidua segregated with fetal sex, with immunologic differences largely contributing at the maternal-fetal interface, maternal response to placentation may also control sexual dimorphism in pregnancy outcomes, for mother and fetus. Future studies will be necessary to determine the specific signaling pathways, and the impact of hormonal regulation on immunologic response that affect sexually dimorphic outcomes.

## Author Contributions

Conceptualization MDP, BRS, HRT, SAK, CRF, SSR, ETW; Data Curation ND, JT, YW, CY, AFK, SDT, CRF; Formal Analysis ND, TLG, TS, AFK, SDT, CRF, MDP; Funding Acquisition MDP, JW, SSR; Investigation TS, TLG, NT, BL, JT, YW, BRS, CY, AFK, SDT, CRF; Methodology TS, TLG, ND, RD, ELC, JT, YW, BRS, CY, AFK, SDT, CRF; Project Administration MDP, TS, TLG; Resources BL, JW; Software ND, JT, YW, AFK, SDT Supervision MDP, SSR; Validation, Visualization ND, TLG, YW, AFK, SDT, MDP; Writing – Original Draft TS, TLG, ND; Writing – Review & Editing TS, TLG, MDP, BRS, HRT, SAK, CRF, SSR.

## Acknowledgments

The authors would like to thank Rae Buttle, Erica Sauro, and Yayu Lin for patient recruitment and sample collection. We gratefully acknowledge the support from the National Institutes of Health, the Eunice Kennedy Shriver National Institute of Child Health and Human Development R01 grants to MDP (R01HD074368, R01HD091773) and the Ruth L. Kirschstein National Research Service Award to TLG (T32DK007770). The content is solely the responsibility of the authors and does not necessarily represent the official views of the National Institutes of Health. The funding agency was not involved in the design, analysis, or interpretation of the data reported.

## Experimental Procedures

### Patient recruitment

Subjects with spontaneously conceived singleton pregnancies were recruited into the Cedars Sinai Medical Center (CSMC) Prenatal Biorepository at time of chorionic villus sampling. With informed consent following IRB-approved protocol (Pro00008600), chorionic villi (placental tissues) and matching decidua (if available) at 10-13 weeks of gestational age were obtained that would otherwise be discarded. All pregnancies were healthy and had a normal karyotype.

### Demographics and pregnancy outcomes

All pregnancies were genetically normal and resulted in healthy term live births, 5 females and 5 males. There is no statistically significant difference in birthweight between female and male infants (p=0.47). (**Table S8)**.

### Tissue storage and processing for total RNA-sequencing

Tissue samples (5-15 mg) were stored with 250 μl RNA*later* RNA Stabilization Reagent (QIAGEN, Hilden, Germany) at −80°C in the CSMC Prenatal Repository until further processing. Tissue was processed as previously described [58]. Briefly, tissue was thawed on ice with 600 μl of RLT Plus lysis buffer (QIAGEN) and 1% β-mercaptoethanol added to each sample. Tissue was homogenized by passing through increasingly thinner single-use needles (22G, 25G, 27G) attached to a 1 mL sterile, RNase-free syringe. The homogenates were loaded onto AllPrep spin columns and the remainder of the protocol was performed following manufacturer instructions for the AllPrep DNA/RNA Mini Kit (QIAGEN). The RNA elution was used for total RNA-sequencing.

### Total RNA-sequencing of tissue samples

RNA-seq libraries were constructed using Illumina TruSeq Stranded Total RNA LT with Ribo-zero kits (Illumina, San Diego). Libraries were pooled at a 4 nM concentration and an average of 19.01 million 2□×□75 bp paired-end reads per sample were generated with Illumina NextSeq 500 using High Output 150-cycle flow cells. Reads were aligned to the human reference genome (build GRCh38 with Ensembl release version 82) using STAR [59]. The DESeq2 Bioconductor package was used in the R statistical computing environment (http://www.R-project.org/) to normalize count data, estimate dispersion, and fit a negative binomial model for each gene [60-62]. P-values were adjusted for multiple comparisons using the Benjamini-Hochberg False Discovery Rate procedure to generate FDR values. To describe tissue expression, we utilized both the Fragments Per Kilobase of transcript per Million mapped reads (FPKM) value and a Z-score describing the position of a gene’s FPKM value compared to all other genes in a sample. The Z-score for a gene equals the number of standard deviations away from the mean FPKM of all genes (in a sample), such that a gene with Z-score of zero (0) has an FPKM equal to the mean FPKM of all genes (in that sample), and genes with a positive Z-score have an FPKM above the mean (in that sample). To determine which genes were expressed in each tissue type, we applied thresholds of mean FPKM>1 and mean Z-score>0 (n=4 per tissue). Consequently, “expressed genes” describes genes which are consistently expressed in decidua or placental tissue. Our criteria for defining differentially expressed genes (DEGs) is FDR<0.10, FPKM>1, and Z-score>0. This is similar to zFPKM method [63].

### Identification of receptors and ligands

Expressed genes from total RNA-seq were filtered by protein-coding biotype, FDR<0.10, FPKM>1, and Z-score>0. We mined the Human Protein Atlas for annotations on all protein-coding genes, data accessed on September 8, 2017 [9]. We used the *DataFrame.merge* function in the *pandas* Python package to cross-reference the RNA-seq genes via matching Ensembl Gene IDs [64]. The initial list of receptors was a merged list of three sources: (1) All confirmed receptors identified by Human Protein Atlas, including nuclear receptors and G-protein coupled receptors; (2) A manual filtering of genes with “receptor” in the gene description after removing false positives such as “non-receptor type” and manually verifying potential hits such as “receptor-like” using annotations from the UniProt protein knowledgebase and the National Center for Biotechnology Information (NCBI) Gene database [7, 8]. Proteins that bound small molecules or extracellular proteins were added as receptors; and, (3) protein-coding genes that had GO terms “receptor” and “binding” and were also identified by the Human Protein Atlas as membrane proteins.

Similarly, the initial list of candidate ligands was compiled from a merged list of: (1) manually filtered genes with “ligand” in the gene description, (2) genes identified as “predicted secreted” by Human Protein Atlas, and (3) genes identified as “plasma proteins” by Human Protein Atlas. Genes that overlapped between the receptor and ligand categories (e.g. *CRLF1*) were manually sorted into one or the other, with preference toward receptor category.

Differentially expressed receptors and candidate ligands (**Table S2** and **Table S3**) were input into the Human Plasma Membrane Receptome (http://www.receptome.org/) to identify interacting receptors and ligands (output) [65]. To identify which output genes are expressed at the maternal-fetal interface, we cross-referenced the database output with our total RNA-seq data to identify which output genes are expressed in decidua or placenta. If a receptor input resulted in additional differentially expressed receptors or ligands (from Human Plasma Membrane Receptome output and our RNA-seq data), then those genes were added to complete **Table S2** and **Table S3**. Since our goal is to identify cell-cell communication pathways relevant for maternal-fetal crosstalk, we kept receptor-receptor pairs and include these when discussing “receptor-ligand interactions” throughout the text. We defined a receptor-ligand pair as maternal-fetal if one gene was placenta-upregulated and its interacting gene was decidua-upregulated, or vice versa. Duplicate pairs were removed in **Table S4**.

### Circos plots

To visualize receptor-ligand interactions, circos plots were generated using *chordDiagram* function in the R package Circlize 0.4.6 (RRID: SCR_002141) and **Table S4** [62, 66]. Gene inputs (“from” column) and gene outputs (“to” column) are interacting receptors and ligands discovered as described earlier. Gene pairs were manually ranked to bring maternal-fetal interactions to the front. The fold-change heatmap colors were generated with function *colorRamp2* from the Circlize package, using black as the middle color corresponding to 1:1 fold-change [66]. Links are comprised of two superimposed lines. Arrows go from ligand to receptor. Receptor-receptor links have arrows on both ends. For circos plots with both single cell and tissue RNA-seq data, gene location selection preferred cell clusters over decidua categories.

### Single cell RNA-seq: preparation of the first trimester placental cells

Fresh placental tissues of 5∼10mg were placed in collection medium and transferred to lab and kept at 4°C overnight before cell preparation. The collection medium was αMEM with sodium heparin (100,000 units/100ml H_2_O, Sigma), gentamycin (0.5%, Invitrogen) and antibiotic-antimycotic (2%, Invitrogen). The tissue was washed in cold PBS to remove maternal contamination and minced with a sterile scalpel, then mixed with 0.25% trypsin, 300U/ml collagenase and 200μg/ml DNAse I and incubated in a 37°C, 5% CO_2_ incubator for 90 minutes with occasional agitation. After centrifugation at 1200 rpm for 10min, cell pellets were carefully resuspended in Chang medium (Irvine Scientific). To avoid overpopulation of erythrocytes in the captured cell population, 1xRBC lysis buffer was applied to the cell suspension for 15min at room temperature, after which, cells were carefully washed and resuspended in 100μl Chang medium and strained using a 70μm flowmi cell strainer (Bel-Art) immediately before library construction.

### Single cell RNA-seq: library construction and sequencing

Single-cell RNA-seq libraries were prepared per Single Cell 3′ v2 Reagent Kits User Guide (10× Genomics, Pleasanton, California). Single cell suspensions were loaded on a Chromium Controller instrument (10× Genomics) to generate single-cell Gel Bead-In-EMulsions (GEMs). GEM-RT were performed in a Veriti 96-well thermal cycler (Thermo Fisher Scientific, Waltham, MA), following which, GEMs were harvested and the cDNAs were amplified and cleaned up with SPRIselect Reagent Kit. Indexed sequencing libraries were constructed using Chromium Single-Cell 3′ Library Kit for enzymatic fragmentation, end-repair, A-tailing, adapter ligation, ligation cleanup, sample index PCR, and PCR cleanup. The barcoded sequencing libraries were quantified using the KAPA Library Quantification Kit (KAPA Biosystems, Wilmington, MA). Sequencing libraries were loaded on a NovaSeq 6000 (Illumina, San Diego, CA) with a custom sequencing setting (26bp for Read 1 and 91bp for Read 2).

### Single cell RNA-seq: data processing and clustering analyses

Cell Ranger 2.0.0 (10× Genomics) was used to demultiplex reads and to convert raw base call files into fastq format. Reads alignment was performed by using STAR (version 2.5.1) [67] with hg38 transcriptome reference from Gencode 25 annotation, containing all protein-coding and long non-coding RNA genes. Raw expression counts for each gene in all samples were collapsed to unique molecular identifier (UMI) counts using Cell Ranger 2.0.0 (10× Genomics). Data filtration and normalization was performed using the workflow described in [68, 69] with Scran 1.8.4 and Scater 1.8.4 packages [70]. Briefly, we filtered cells with log-library sizes that are more than 2 median absolute deviations (MADs) below median as well as cells with log-transformed expressed genes that are more than 2 MADs below median. Cells with mitochondrial proportions of 2 MADs higher than median were also removed. In addition, the low abundance genes (average UMI counts<0.1) were excluded, yielding 7,245 cells and 6,806 genes in total for further analyses, with median genes of 1,860 per cell and median UMI counts of 5,654 per cell. The data matrix was then cell-specifically normalized by deconvolution method using centered pool-based size factor [69] for similar cells with clustering. We applied a shared nearest neighbor graph-based algorithm to cluster the cells for global population. To obtain two-dimensional visualization of the cell population, principal components analysis was first run on the normalized log-expression values. Top 10 principal components that explained the most variability were selected to perform t-Distributed Stochastic Neighbor Embedding (t-SNE) for cells using default parameters. For tSNE, the perplexity parameter and the _ parameter were set to 30 and 0.5, respectively, while the other parameters were left as default and total iterations was 1000. For sub-clustering analysis within the trophoblast population, an average UMI cutoff of 0.05 was applied to remove low-abundance genes, and normalization using the same method as above was performed. A total of 1,465 cells and 8,214 genes were used, and hierarchical clustering with a dynamic tree cut was performed to define sub-clusters of trophoblast cells. For analyses on both global and trophoblast populations, outliers in tSNE were excluded.

### Single cell RNA-seq: identification of differentially expressed genes (DEGs)

The edgeR 3.22.4 package [71] was applied to conduct differential expression analysis. Briefly, after initial filtration and normalization, the SingleCellExperiment object was converted to a DGEList object containing the UMI counts and calculated size factors. After dispersion estimation using a design matrix, differentially expressed genes were identified between any two cell types by fitting a negative binomial generalized log-linear model to the counts for each gene. The identification of DEGs between any two cell types was adjusted by samples. Benjamini-Hochberg method was applied for multiple test corrections to calculate false discovery rates (FDR). Pairwise comparison between cell types was performed, and the DEGs (Log_2_FC>1, FDR<0.01) that were specifically enriched for each cell type were depicted as a heatmap (genes that are expressed in at least 5% of cells from all conceptions were selected). For DEG analyses between sexes in each cell type, the DE genes (FDR<0.01) from both male and female conceptions were selected for enrichment analysis. A stricter cutoff of FDR<0.01 and |Log_2_FC|>1 was applied to highlight the most significant genes between sexes.

### Enrichment analysis and upstream regulator analysis

For enrichment analysis in tissue: Receptors and ligands significantly upregulated in placenta or decidua (FDR<0.10, FPKM>1, Z-score>0) were used for core analyses with Ingenuity Pathway Analysis software (RRID: SCR_008653). Inputs used were Ensembl Gene IDs, FDR value, and log_2_ fold-change from DESeq2 [61]. Protein-coding genes upregulated in placenta and decidua were also analyzed, with similar results.

For single cell analysis: The Gene Ontology Consortium website (http://www.geneontology.org/) revealed Enriched Gene Ontology terms (biological processes) using the cluster-specific genes. Upregulated genes were used for enrichment analysis for fetal sexes (FDR<0.01) or comparing two sub-clusters of trophoblast cells (Log_2_FC>1, FDR<0.01). Bonferroni-corrected for P<0.05 was used for multiple testing. For upstream regulator analyses using Ingenuity Pathway Analysis (RRID: SCR_008653), input was significantly and specifically expressed genes in each cell type following DEG analysis and pairwise comparison. Molecule types of upstream regulators included were: endogenous mammalian chemicals, cytokines, enzymes, G-protein coupled receptor, growth factor, ion channel, kinase, ligand-dependent nuclear receptor, mature miRNA, miRNA, peptidase, phosphatase, transcription regulator, translation regulator, transmembrane receptor, transporter.

### Pseudotime analysis

To delineate the developmental trajectory of the trophoblast cells, previous sub-clusters within trophoblast cells were applied using Monocle 2.8.0 [72-74]. Cells were ordered in a semi-supervised manner by using marker genes in each sub-cluster within the CellTypeHierchy system.

## Supplemental item titles and legends

**Figure S1. Principal components analysis of RNA-seq results.** Plot shows clustering of tissue samples, verifying sample purity. CVS = chorionic villus sampling (placenta/fetal contribution). DEC = decidua (maternal). M1, M2 = pregnancy with male fetus. F1, F2 = pregnancy with female fetus.

**Figure S2. Venn diagram for maternal-fetal interface DEGs from total RNA-sequencing. (A)** Differentially expressed protein-coding genes. **(B)** Differentially expressed receptors. **(C)** Differentially expressed ligands. Expressed genes reach thresholds of FPKM>1 and Z-score>0 in decidua, placenta, or both. Up arrow indicates direction of significant upregulation (FDR<0.10).

**Figure S3. Circos plot with “DEG-NS” receptor interactions.** Circos plot for 211 “DEG-NS” interactions (one gene is FDR<0.10, but the other is not significantly different). Links are colored by direction of upregulation: green if both genes directionally up in placenta (fetal-fetal), pink if both genes are directionally up in decidua (maternal-maternal), black if directions of upregulation are opposite (maternal-fetal). One link is colored yellow because the not significant gene (FDR>0.10) has a mismatch in tissue-uniqueness and upregulation direction. Tracks are similar to **Figure 1A**.

**Figure S4. Sexual dimorphism of enriched GO terms (in additional cell types)** The top 10 enriched GO terms are presented as −log_10_(P value) for either sex in cell types in addition to trophoblasts. The genes upregulated in either male or female conceptions (Log_2_FC>0) were input in gene ontology consortium (http://geneontology.org/) for enriched GO biological processes (Bonferroni-corrected for P < 0.05 was used for multiple testing).

**Table S1. Differentially expressed protein-coding genes in tissue.** DESeq2 identified 3811 genes significantly different (FDR<0.10) between decidua and placental tissue.

**Table S2. All significantly differentially expressed receptors.** Includes 356 receptors with FDR<0.10.

**Table S3. All significantly differentially expressed ligands.** Includes 1043 candidate ligands with FDR<0.10.

**Table S4. Interacting gene pairs at the first trimester maternal-fetal interface**. Includes 337 gene pairs where at least one gene is FDR<0.10.

**Table S5. Placental cell markers** Cluster-specific markers emerged from pairwise comparison among the clusters are ranked by normalized expression UMI counts from the most abundant to the least (log_2_FC>1, FDR<0.01).

**Table S6. Trophoblast subcluster markers** Markers specifically expressed in sub-clusters of trophoblast cells (log_2_FC>1, FDR<0.01), selected from pairwise comparison among the seven sub-clusters are ranked by normalized UMI counts from the most abundant to the least. No markers emerged for sub-cluster 1 and 7 from pairwise comparison, therefore markers of sub-clusters 2, 3, 4, 5, 6 are included in the table.

**Table S7. Placental cell type-specific sexually dimorphic genes** Fold changes of top sexually dimorphic genes (ILog_2_FCI>1, FDR<0.01, shown in bold) in each cluster are presented in table S7A (upregulated in cells from male conceptions) and S7B (upregulated from female conceptions). The significantly sexually dimorphic expression patterns (FDR<0.01) of these genes, although may not meet the criteria of ILog_2_FCI>1 in some clusters, are shown for all clusters. * indicates the genes that are expressed in less than 5% of cells (excluded in Figure 5B).

**Table S8. Patient demographics and pregnancy outcomes** Demographic and pregnancy outcome information is shown for all 10 patients, including 4 patients in total RNA-seq analysis and 6 patients in single cell RNA-seq analysis.

