## Supplementary figures and images for "The maternal-fetal interface of successful pregnancies and impact of fetal sex using single cell sequencing"

### Supplemental Figure 1

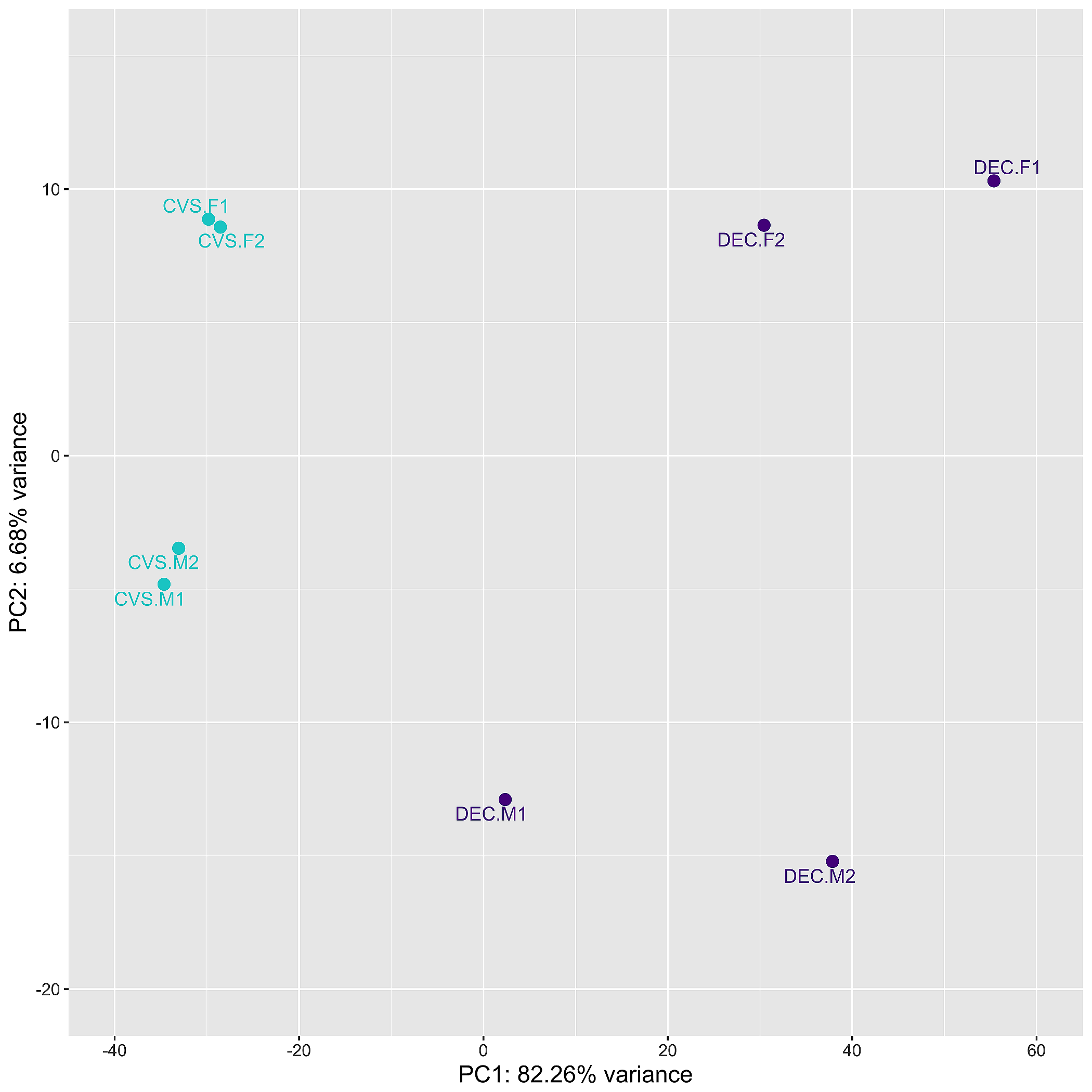

### Supplemental Figure 2

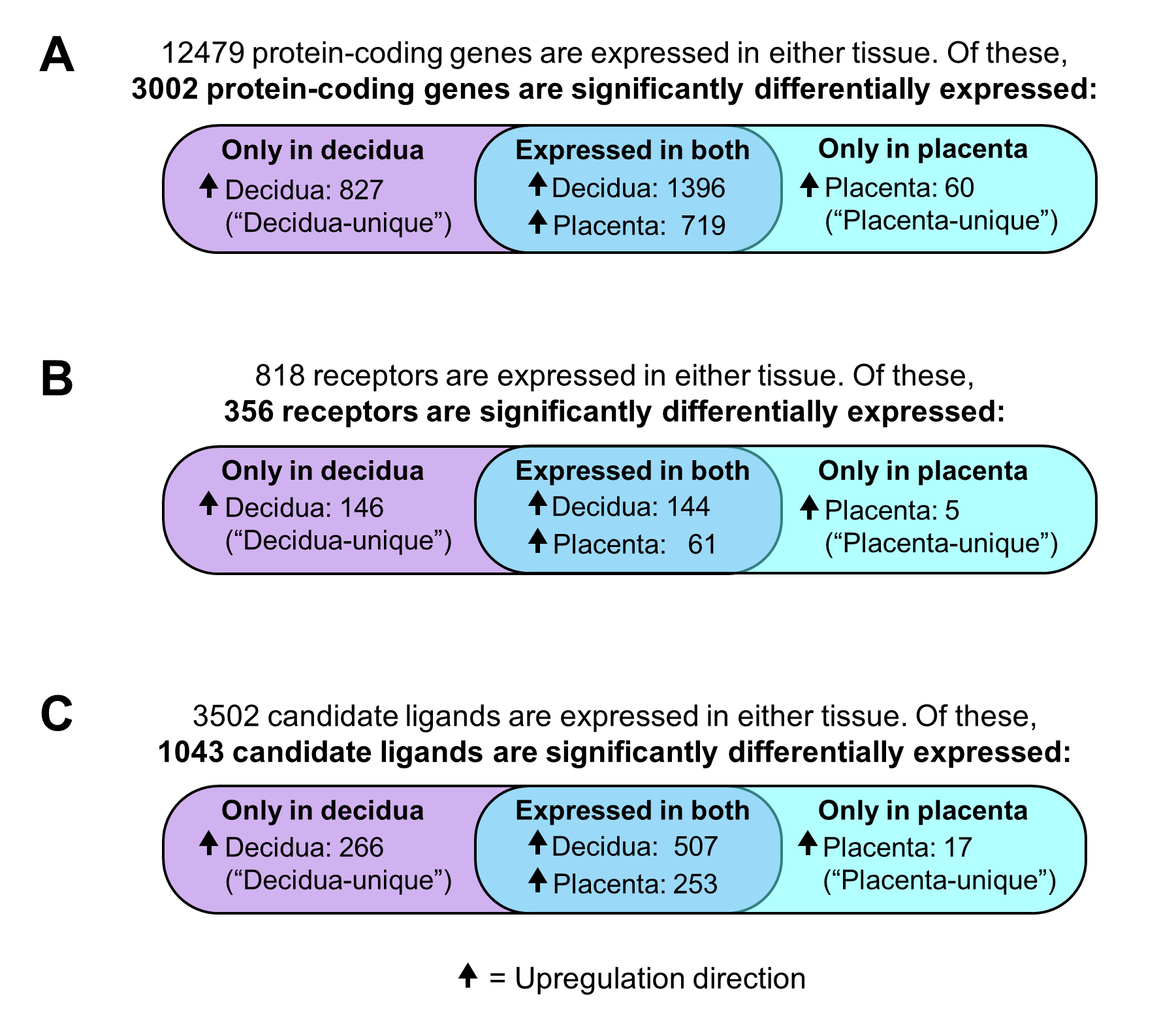

### Supplemental Figure 3

**Supplemental Figure 3. Circos plot with 211 DEG-NS receptor interactions (264 genes total).**

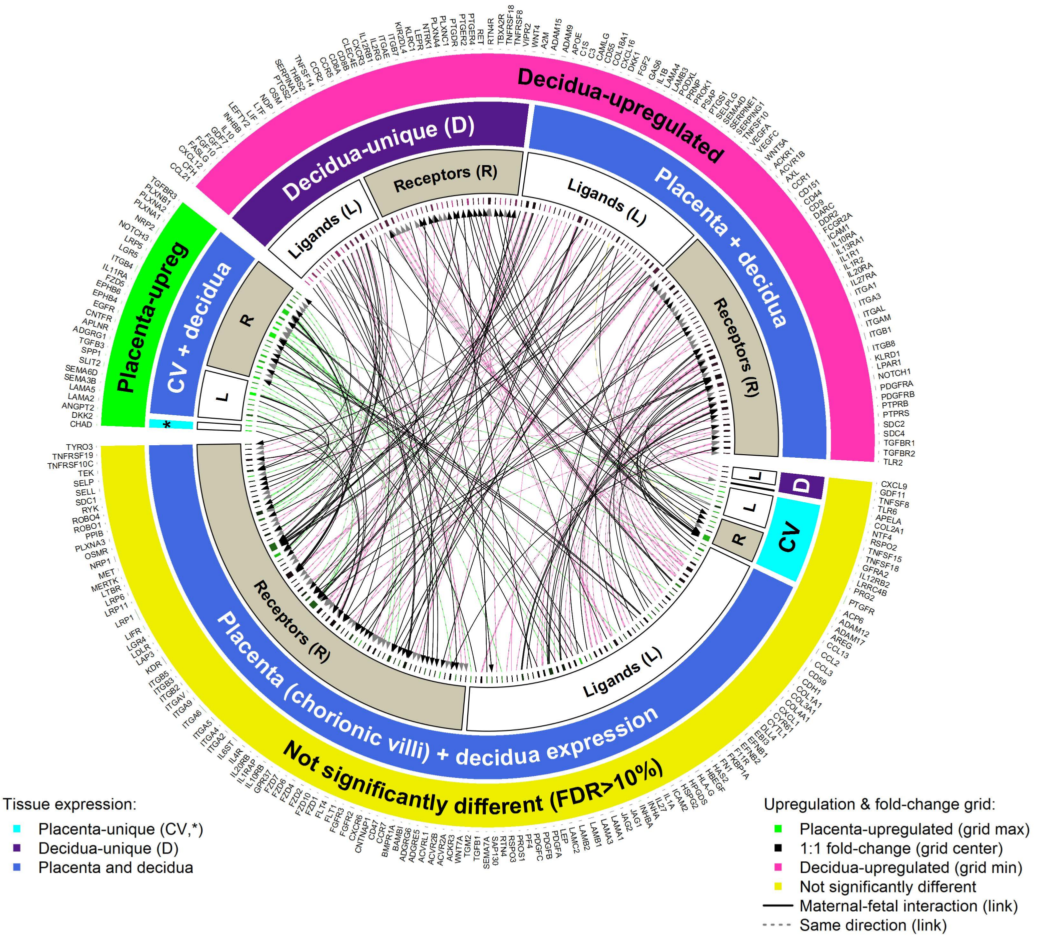

### Supplemental Figure 4

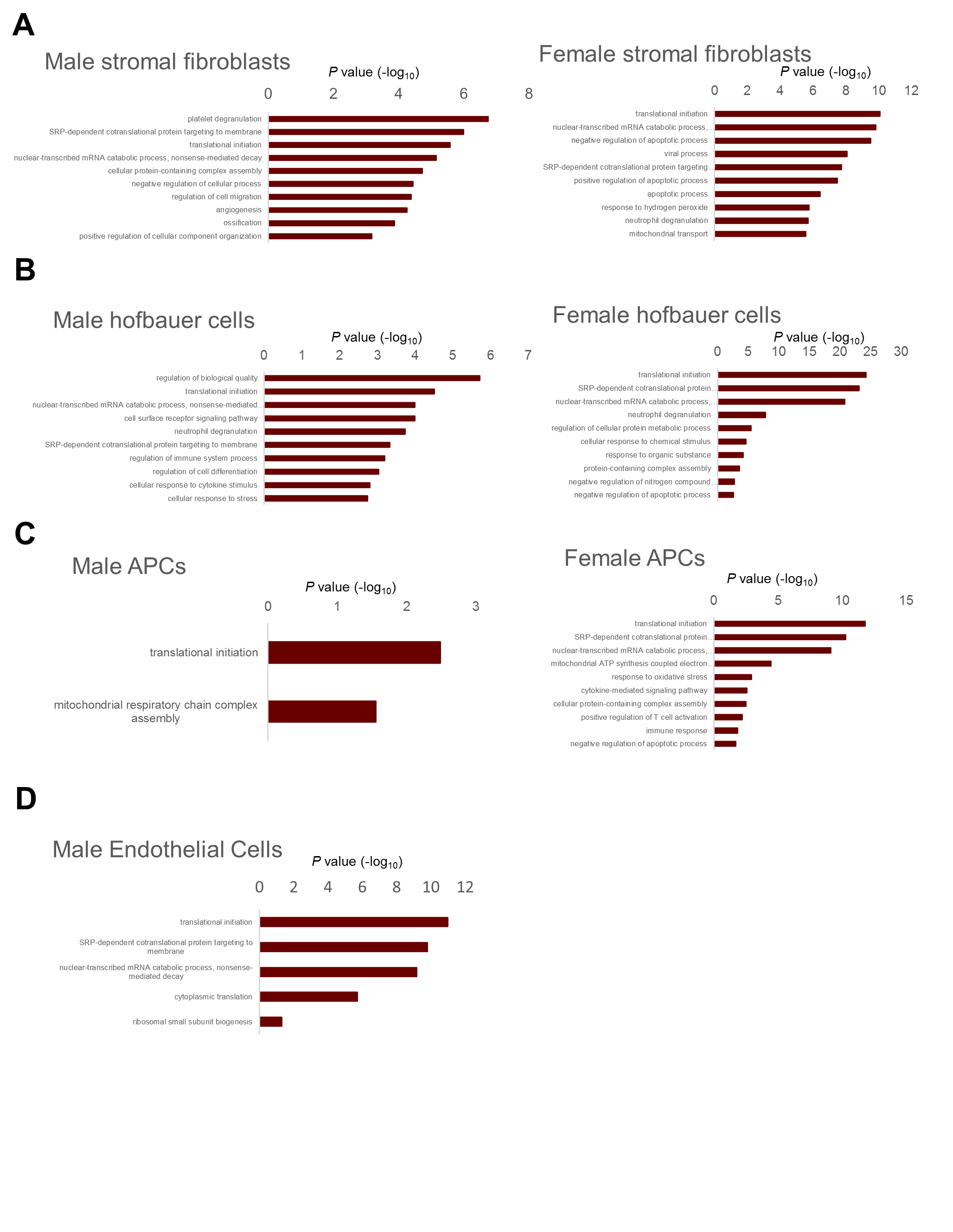
