## Supplemental Table 7 for "The maternal-fetal interface of successful pregnancies and impact of fetal sex using single cell sequencing"

Table S7A. Top genes upregulated in placental cells from male conceptions

| **Gene** | **Chr** | **Biotype** | **Trophoblast Cells** | **Stromal Cells** | **Hofbauer Cells** | **APCs** | **Endothelial Cells** |
| --- | --- | --- | --- | --- | --- | --- | --- |
| **DDX3Y** | Y | Protein | **3.58** | **4.14** | **6.79** |  |  |
| **EIF1AY** | Y | Protein | **8.38** | **4.08** | **2.53** |  | **5.41** |
| **RPS4Y1** | Y | Protein | **23.65** | **45.51** | **21.08** | **2.21** | **20.26** |
| **RPL36A** | X | Protein | 1.65 | 1.78 | 1.59 | 1.70 | **2.49** |
| **HLA-C** | 6 | Protein | **2.36** | 1.35 |  |  |  |
| **MUC15** | 11 | Protein | **2.21** |  |  |  |  |
| **NOTUM** | 17 | Protein | **3.24** |  |  |  |  |
| **SNHG19** | 16 | lincRNA | **2.01** | 1.33 | 1.32 |  |  |
| **SNHG25** | 17 | lincRNA | **2.03** | 1.66 | 1.24 |  |  |
| **BCYRN1** | 2 | scRNA |  | **2.85** |  |  |  |
| **CTC-425F1.4** | 19 | antisense | 1.94 | **2.32** | 1.69 |  |  |
| **HLA-DRB1** | 6 | Protein |  | **2.12** |  |  |  |
| **STC1** | 8 | Protein |  | **2.14** |  |  |  |
| **APOC1** | 19 | Protein | 1.49 | 1.93 | **2.02** | 1.38 |  |
| **BDH1** | 3 | Protein | 1.43 | 1.79 | **2.19** | 1.93 |  |
| **FCGBP** | 19 | Protein |  | 1.10 | **2.41** |  |  |
| **DONSON** | 21 | Protein | 1.50 | 1.94 | **3.36** | **2.77** | **5.17** |
| **MT1G*** | 16 | Protein |  |  | **3.91** | **4.36** |  |
| **PRDM6** | 5 | Protein | 1.40 | 1.37 | **2.10** | **2.38** |  |
| **TMEM176A** | 7 | Protein |  | 1.47 | **2.19** | **2.82** |  |
| **TMEM176B** | 7 | Protein |  | 1.76 | **2.33** | **3.58** |  |
| **BEST1** | 11 | Protein |  |  | 1.45 | **2.39** |  |
| **CH17-373J23.1** | 1 | LincRNA |  | 1.48 |  | **2.21** |  |
| **FBLIM1** | 1 | Protein | 1.41 | 1.37 | 1.28 | **2.09** |  |
| **IL1RN** | 2 | Protein |  |  |  | **2.93** |  |
| **MMP9** | 20 | Protein |  |  |  | **2.54** |  |
| **NBEAL1** | 2 | Protein | 1.71 | 1.95 | 1.54 | **2.02** |  |
| **HLA-A** | 6 | Protein |  | 1.35 | 1.50 |  | **2.56** |
| **IGF1** | 12 | Protein |  |  |  |  | **2.27** |
| **RPS21** | 20 | Protein | 1.24 | 1.44 | 1.35 | 1.42 | **2.16** |
| **TXNIP** | 1 | Protein |  |  |  |  | **2.76** |

Table S7B. Top genes upregulated in placental cells from female conceptions

| **Gene** | **Chr** | **Biotype** | **Trophoblast Cells** | **Stromal Cells** | **Hofbauer Cells** | **APCs** | **Endothelial Cells** |
| --- | --- | --- | --- | --- | --- | --- | --- |
| **MAGEA4** | X | Protein | **6.72** | 1.12 |  |  |  |
| **TMSB4X** | X | Protein | **2.00** |  |  | 1.20 |  |
| **XIST** | X | LincRNA | **8.93** | **13.32** | **7.71** | 1.53 | **9.94** |
| **CCL3** | 17 | Protein | **2.32** | 1.30 |  |  |  |
| **CCL4** | 17 | Protein | **2.25** |  |  |  |  |
| **GTSF1** | 12 | Protein | **2.24** | 1.80 |  |  |  |
| **RNASE1** | 14 | Protein | **2.06** |  |  | 1.60 |  |
| **CXCL8** | 4 | Protein | **2.05** | 1.32 |  | 1.57 |  |
| **HBG2** | 11 | Protein | 1.89 | **3.75** | 1.59 |  | **13.92** |
| **GSTT2B** | 22 | Protein |  | **2.43** | 1.18 |  |  |
| **SNHG7** | 9 | LincRNA |  | **2.30** | 1.82 |  |  |
| **HBB** | 11 | Protein | 1.60 | **2.16** | **3.55** | 1.57 |  |
| **RGS1** | 1 | Protein |  |  | **3.26** |  |  |
| **CCL13** | 17 | Protein |  |  | **2.13** |  |  |
| **RETN** | 19 | Protein |  | 1.18 | **2.07** |  |  |
| **GPR183** | 13 | Protein | 1.27 |  | **2.33** | **2.78** |  |
| **CGA** | 6 | Protein | 1.58 | 1.77 | **2.11** | **3.98** |  |
| **F13A1** | 6 | Protein | 1.64 |  | 0.89 | **2.10** |  |
| **MTRNR2L1** | 17 | Protein | 0.57 |  | 1.26 | **2.66** | **3.16** |
| **GADD45G** | 9 | Protein | 1.66 | 1.17 |  | 0.66 | **3.83** |
